# Disease-associated mutations directionally modulate epigenetic age in iPSC-derived tissues

**DOI:** 10.64898/2026.09.30.755445

**Authors:** Nicholas J. Skvir, Todd W. Dowrey, Samuel F. Cranston, Elise MacDonald, Richard Giadone, Dakota Jones, Carly S. Golden, Braden Pate, Madeline Labott, Maria A. Serrano, Konstantinos-Dionysios Alysandratos, Kim Vanuytsel, George J. Murphy

## Abstract

Reprogramming somatic cells into induced pluripotent stem cells (iPSCs) resets epigenetic age, creating a major challenge for modeling age-associated cellular states and limiting the use of iPSCs in aging research beyond the genetic component. Recent work has shown that healthy organoids can re-accumulate age-associated epigenetic states over extended culture; whether disease-associated genetic states can directionally reshape these trajectories after reprogramming remains unknown. Here, we present a reproducible, *in-vitro* approach to re-establish epigenetic aging trajectories using iPSC-derived organoid systems, as quantified by multiple epigenetic clock models. We first demonstrate that epigenetic age progressively increases with serial passaging in a lung organoid model (alveolospheres). We then show that disease-associated mutations in multiple models drive additional, cell-intrinsic shifts in epigenetic age, implicating disease state as an independent modulator of epigenetic clocks. Together, these findings reinforce that iPSC-derived organoids recapitulate epigenetic aging trajectories over time, while highlighting disease state as a potent and potentially tractable modulator of epigenetic age *in vitro*. By enabling more faithful modeling of age-associated cellular states, this approach expands the utility of iPSC-based systems for dissecting the molecular mechanisms that regulate human aging.

## Introduction

The advent of induced pluripotent stem cell (iPSC) technology has opened new avenues for studying human disease *in vitro*, enabling the generation of patient-specific cell lines that capture individual genetic backgrounds. This may be particularly valuable in the context of aging research, where access to primary aged human tissue (i.e. brain) is limited and difficult to standardize. However, a fundamental limitation of this approach comes from the reprogramming process itself. It is well known that the reprogramming process resets the epigenetic landscape of somatic cells to a pluripotent state^1,2^, effectively erasing molecular signatures of age that were present in the donor tissue^3,4^. Although epigenetic rejuvenation is a remarkable feature of iPSC generation, and has spurred substantial interest in partial reprogramming^4–6^, it presents a significant challenge for the development of iPSC-based models of aging, as reprogramming erases epigenetic information that may contribute causally to the process^7–12^.

Epigenetic clocks offer a powerful framework for estimating biological age at the molecular level^3,13^. These computational models leverage DNA methylation patterns at specific CpG sites to predict chronological or biological age across numerous tissues^14–16^ and species^17^. These clock models are calibrated to capture distinct aspects of the aging process, with some optimized explicitly for tracking chronological age^18^, and others tuned to relate to changes in morbidity and mortality risk^19,20^. In iPSCs and iPSC-derived cells however, these clocks consistently report ages near zero^3,21–23^ following reprogramming, and few studies have systematically examined whether epigenetic age can be re-instilled in this context.

Organoids represent a conceptually distinct context from conventional monolayer iPSC-derived cultures. Their three-dimensional architecture, cellular heterogeneity, and capacity for extended culture introduce biological complexity that may more faithfully recapitulate aspects of *in vivo* tissue development^24,25^ and aging^26,27^. Whether this complexity creates conditions permissive to epigenetic age accumulation, and whether disease-associated mutations might further modulate such trajectories in biologically meaningful ways, has only recently begun to be addressed. Work in human brain organoids has shown that cultures maintained for up to five years accumulate clock-predicted epigenetic age that tracks time in culture and parallels *in vivo* brain maturation^28^, establishing that epigenetic age can re-emerge after reprogramming, albeit on a human developmental timescale. Whether epigenetic age can accumulate in other organoid lineages, and at what rate, remains largely unexplored. The effect of disease on epigenetic age, by contrast, has been examined primarily *in vivo*, where monogenic disease mutations in patients have been associated with shifts in blood-based epigenetic age^29,30^. Yet whether these effects are cell-intrinsic or depend upon the systemic disease environment cannot be determined from patient samples alone, and is a question that isogenic iPSC-derived models can test directly.

Here we show that iPSC-derived lung organoids (alveolospheres) accumulate epigenetic age in a time-dependent and reproducible manner, and that disease-associated mutations in these lines, as well as in two additional disease models in different tissues, shift epigenetic age in directions consistent with the known biology of each condition. Using established DNA methylation clocks and transcriptional profiling, we studied multiple iPSC-derived organoid and monolayer models across timepoints and disease conditions, finding that clock predictions diverge in informative ways depending on the nature of the disease perturbation. Together, these results suggest that controlled induction of epigenetic age in iPSC-derived tissues is an achievable goal, and that disease-relevant mutations represent one avenue toward this end. If such induction can be further refined and generalized, iPSC-based models stand to gain considerable power as platforms for studying the epigenetic underpinnings of aging.

## Results

### Epigenetic age is reset during iPSC reprogramming across multiple clock models

To establish a baseline, and to confirm that the epigenetic age reset associated with iPSC reprogramming was reproducible in our hands, we profiled peripheral blood mononuclear cells (PBMCs) from a 54-year-old donor across a reprogramming time course, sampling at baseline Day 0 (prior to Sendai viral transduction), Days 1, 3, 9, and 13 post-transduction, and then at clonal passages 1 and 3. Epigenetic age was assessed using four clock models: Horvath^3^, Hannum^13^, EnCen40+^31^, and PhenoAge^19^. Across all four models, predicted epigenetic age declined gradually before dropping more sharply after approximately D9, converging toward near-zero values by the clonal passage stage (**Figure 1A**). This pattern is consistent with the known kinetics of reprogramming, in which pluripotency emerges over the course of epigenetic remodeling rather than immediately following transduction^32,33^. This reset establishes the baseline against which the elevated epigenetic ages observed in our organoid models are interpreted.

**Figure 1.**
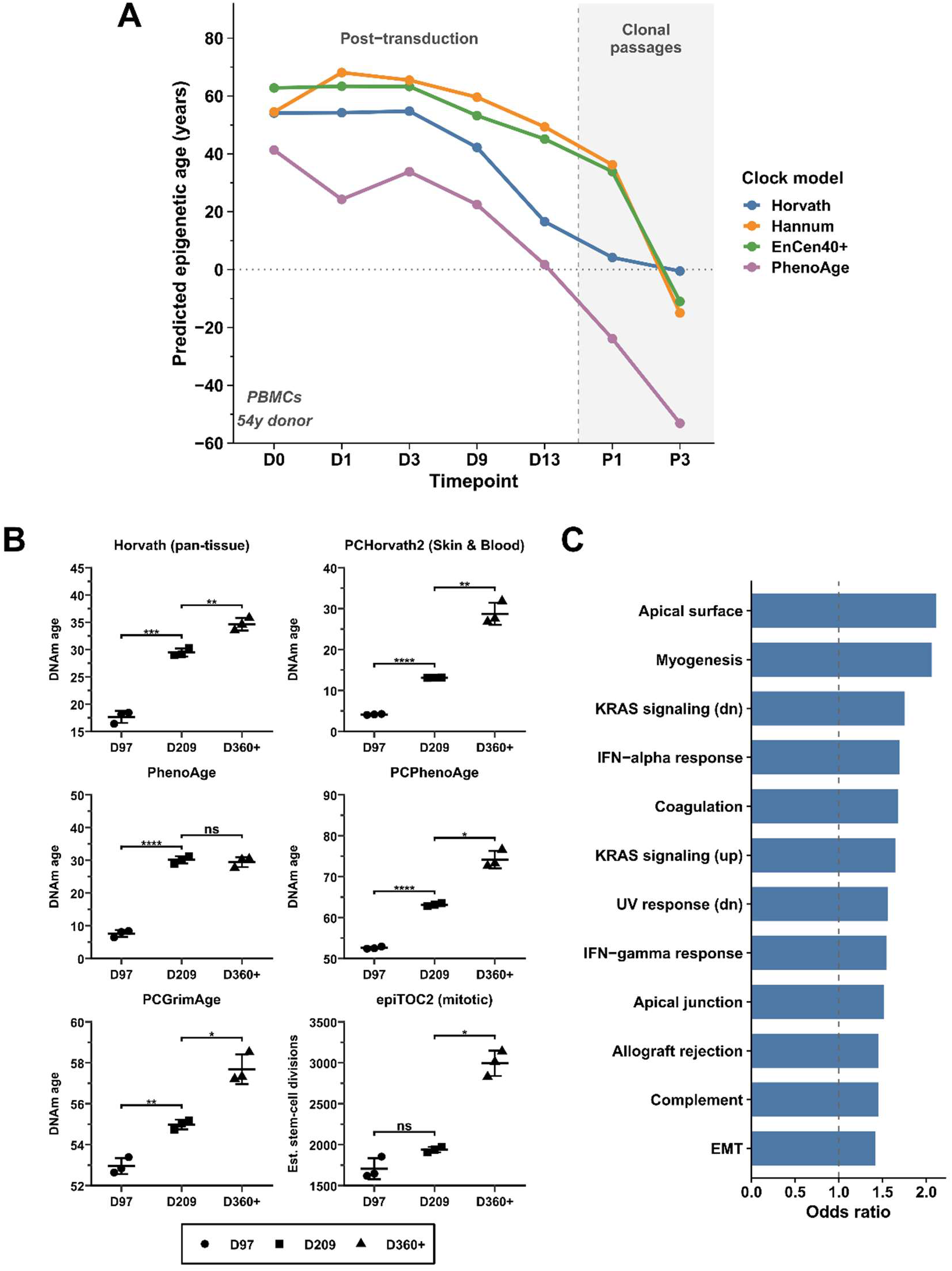
Epigenetic age is reset during iPSC reprogramming and re-accumulates in alveolosphere organoids over extended culture. **(A)** Predicted epigenetic age of peripheral blood mononuclear cells (PBMCs) derived from a 54-year-old donor profiled across an iPSC reprogramming time course. Samples were collected at baseline (D0), post-transduction timepoints (D1, D3, D9, D13), and clonal passages (P1, P3). Four epigenetic clock models are shown: Horvath, Hannum, EnCen40+, and PhenoAge. Shaded region indicates clonal passage stage. Note that PhenoAge is a linear model with no lower bound; negative values indicate an epigenetically young state and should be interpreted in relative rather than absolute terms. **(B)** Predicted epigenetic age of SFTPC-corrected iPSC-derived alveolosphere organoids at three culture timepoints (D97, D209, D360+), assessed by six DNA methylation clocks: the Horvath pan-tissue clock, PhenoAge, PC PhenoAge, PC Skin & Blood, PCGrimAge, and the epiTOC2 mitotic clock, which estimates cumulative stem-cell divisions. Each clock is plotted on its own axis because the clocks differ in units and scale. Individual data points are shown with mean ± SD (n=3 per timepoint). Welch’s t-test with Holm-Šídák correction were performed for each model (* p<0.05, ** p<0.01, *** p<0.001, **** p<0.0001, ns = not significant). **(C)** Over-representation analysis (ORA) of DMR-proximal genes in control alveolospheres with passaging, comparing the early passaged samples (D97) to the late passaged sample (D360+). Bars represent odds ratio for each significantly enriched term (FDR < 0.05). Analysis performed using EnrichR against the MSigDB Hallmark 2020 gene set database.

### Epigenetic age accumulates progressively in iPSC-derived alveolosphere organoids over extended culture

To investigate whether iPSC-derived organoids could accumulate epigenetic age beyond the near-zero values typically observed in standard iPSC-derived cultures^22^, we profiled the DNA methylation-based epigenetic age of iPSC-derived alveolosphere organoids^34,35^. Organoids were assessed in biological triplicate at three timepoints spanning early to late culture: D97, D209, and two later timepoints (two replicates at D360 and one at D417). Predicted epigenetic age across all models profiled increased steadily and monotonically with time in culture, with low inter-replicate variance (**Figure 1B**). These findings demonstrate that iPSC-derived alveolosphere organoids accumulate epigenetic age in a time-dependent and reproducible manner, reaching predicted ages of nearly 30 years by late culture (as measured by the Horvath clocks and PhenoAge) and substantially exceeding those reported for conventional iPSC-derived monolayer cultures^22^. This accumulation also notably outpaces time in culture, contrasting with previous findings in which the epigenetic age of brain organoids closely tracks calendar time^28^.

### Extended culture induces global hypomethylation and aging-associated pathway remodeling

To determine whether the progressive increase in clock-predicted age reflected broader changes across the methylome, we examined genome-wide methylation distributions and differentially methylated regions (DMRs) in alveolospheres across culture time. With extended culture, the proportion of highly methylated CpGs progressively decreased, accompanied by a corresponding increase in unmethylated CpGs (**Supplemental Figure S1A**). This shift toward global hypomethylation parallels an established hallmark of *in vivo* aging, in which the genome undergoes progressive loss of DNA methylation at repetitive elements, intergenic regions, and gene bodies over time^36,37^, and provides orthogonal support for the aging trajectories detected by the epigenetic clocks.

The early-versus late-passage comparison yielded 8,923 unique DMR-proximal genes, the largest gene set identified across the comparisons examined. Over-representation analysis (ORA) revealed significant enrichment (FDR < 0.05) of immune and inflammatory pathways, including Interferon Gamma and Alpha Response, Complement, Coagulation, and Allograft Rejection, together with Epithelial Mesenchymal Transition and KRAS Signaling; Hedgehog Signaling was nominally enriched (FDR = 0.084) (**Figure 1C; Supplemental Table 1**). The immune and inflammatory terms are consistent with the chronic low-grade inflammation associated with aging, often referred to as inflammaging^38,39^, while the enrichment of EMT-associated genes is consistent with previous descriptions of fibrotic disease contexts^40,41^. Together, these genome-wide methylation and pathway data support the interpretation that increased clock estimates reflect a biologically meaningful aging-like state.

### SPC2 mutant alveolospheres show altered epigenetic age profiles relative to isogenic controls

Having established that alveolosphere organoids accumulate epigenetic age over extended culture, we next asked whether a disease-associated mutation would further modulate this trajectory. We profiled isogenic alveolosphere lines carrying a mutation in SFTPC, which encodes surfactant protein C and is expressed in alveolar epithelial type II cells (AEC2s)^42^. This mutation has been previously characterized as causing proteostatic stress, inflammation, impaired autophagy, and metabolic dysregulation, phenotypes that overlap substantially with canonical hallmarks of aging^36^ and make these mutants compelling candidates for epigenetic age acceleration. These lines derive from the previously characterized SFTPC^I73T^-mutant iPSC line (hereafter “SPC2”; see methods). Mutant and corrected isogenic lines were profiled at D97 and D209 using the PhenoAge model and the Horvath skin & blood clock. As a second generation clock^43^, PhenoAge is calibrated to reflect biological age in relation to morbidity and mortality risk, and may therefore be more sensitive to the pro-inflammatory and proteostatic perturbations characteristic of the mutation, while the skin and blood clock is specifically well-calibrated to cells in culture. PhenoAge revealed a robust and consistent elevation in epigenetic age in SPC2 mutants relative to corrected controls, of approximately 29 years at D97 and 27 years at D209, while the Horvath skin and blood clock showed relatively smaller yet still significant elevation at both time points (**Figure 2A**). Additionally, previously published data with these models indicates that SPC2 mutant alveolospheres exhibit reduced replication relative to corrected controls, arguing against increased replicative burden as an explanation for the elevated PhenoAge signal^42^. These results are also borne out by the predictions made by epiTOC2^44^, a mitotic clock trained to predict cumulative cell divisions. The concordance between the known disease phenotype and the directions of these signals suggests that this clock may be capturing biologically meaningful epigenetic dysregulation in this context.

**Figure 2.**
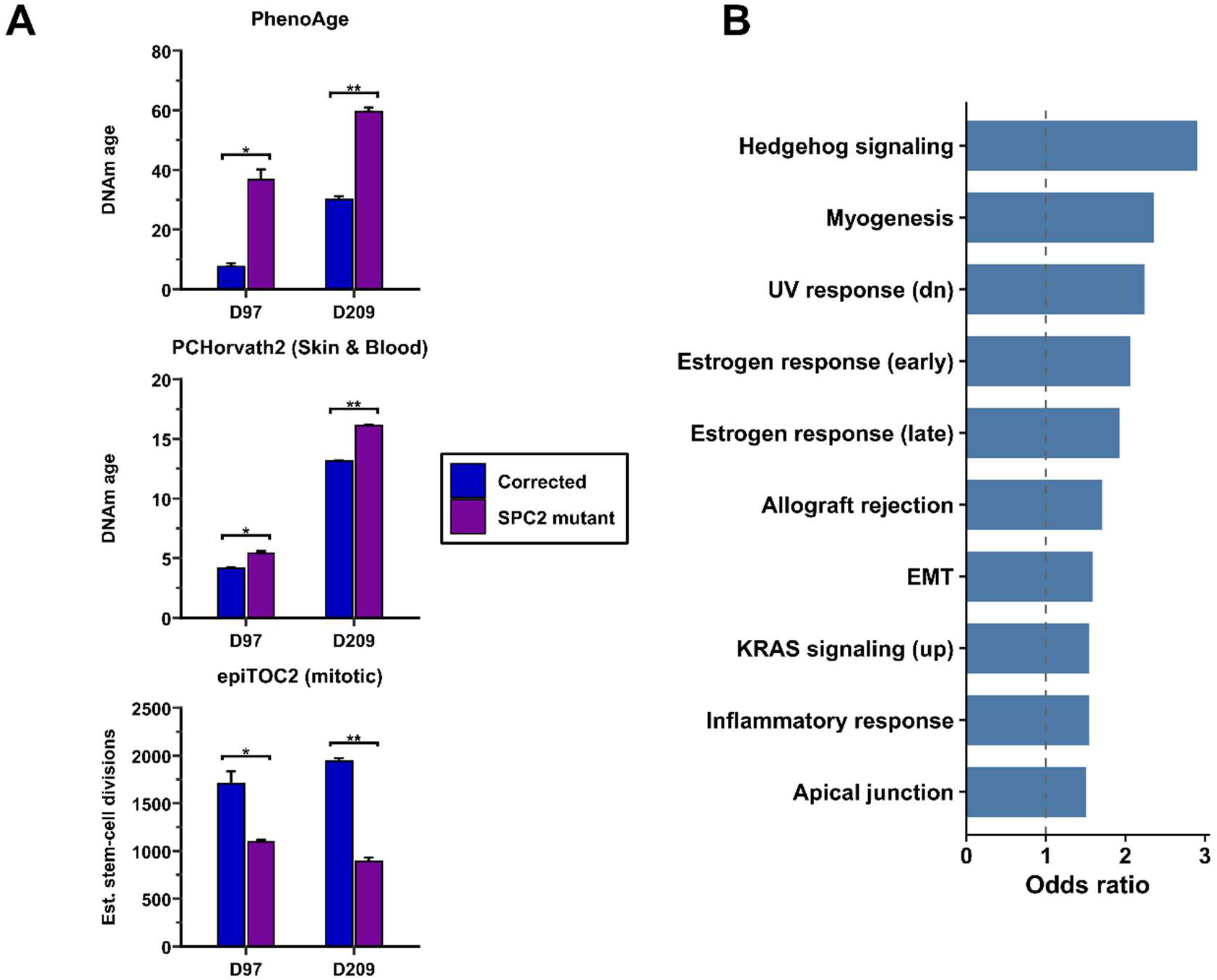
SPC2 mutant alveolospheres show altered epigenetic age profiles relative to isogenic corrected controls. **(A)** Predicted epigenetic age of corrected and SFTPC-I73T (SPC2) mutant alveolosphere organoids at D97 and D209, estimated with three DNA methylation clocks: PhenoAge, PCHorvath2 (Skin & Blood clock), and the epiTOC2 mitotic clock, which estimates cumulative stem-cell divisions rather than age in years. Bar graphs show mean ± SD. Corrected: n=3; SPC2 mutant: n=2 per timepoint. Mutant and corrected organoids were compared within each timepoint by Welch’s t test, with Holm-Šídák correction (* p<0.05, ** p<0.01). Mutant organoids were epigenetically older by these models while showing fewer estimated stem-cell divisions. **(B)** Over-representation analysis (ORA) of DMR-proximal genes in SPC2 mutant versus corrected alveolospheres. Bars represent odds ratio for each significantly enriched term (FDR < 0.05). Analysis performed using EnrichR against the MSigDB Hallmark gene set database.

### SPC2 mutation drives aging-associated epigenetic remodeling with a distinct inflammatory program

Interestingly, the methylation landscape of SPC2 mutant alveolospheres differed from that observed during extended culture. Although the highly methylated CpG peak was reduced relative to corrected controls, methylation values were redistributed primarily toward intermediate levels rather than toward the broad hypomethylated state observed with passaging (**Supplemental Figure S1B**). This pattern suggests that culture duration and the SPC2 mutation alter the methylome through partially distinct mechanisms. The SPC2 mutant versus corrected comparison identified 5,625 unique DMR-proximal genes. Significantly enriched terms included Estrogen Response, UV Response, Allograft Rejection, Hedgehog Signaling, Epithelial Mesenchymal Transition, Inflammatory Response, and KRAS Signaling (**Figure 2B; Supplemental Table 2**). Several terms, including Myogenesis, EMT, Hedgehog Signaling, and Apical Junction, were also enriched in the passaging comparison, indicating partial convergence between disease-associated remodeling and the aging-like changes induced by extended culture. Inflammatory Response, however, reached significance specifically in the SPC2 comparison, consistent with the previously characterized pro-inflammatory phenotype of this mutation^42^, and was not among the significant hits in the passaging comparison. The SPC2 mutation therefore does not simply reproduce the methylation changes caused by extended culture. Instead, it appears to combine features of aging-associated epigenetic remodeling with a distinct inflammatory program reflective of the underlying disease biology, consistent with the marked elevation in PhenoAge observed in SPC2 mutant organoids.

### KMT2D mutant brain organoids show epigenetic signatures consistent with developmental arrest

To examine whether disease-associated epigenetic dysregulation could be detected in a brain organoid model, we profiled isogenic cortical brain organoids carrying a loss-of-function mutation in KMT2D, which causes Kabuki Syndrome^45,46^. KMT2D encodes a histone methyltransferase required for the activation of developmental enhancers, and its loss has been shown to disrupt the coordination of core lineage transcription factors, cause accumulation of neural progenitors in G1 phase following neural induction, and impair the resolution of stable cell lineages during early neurodevelopment46. Because KMT2D directly shapes the enhancer chromatin landscape, its loss represents a primary perturbation of the epigenetic machinery interrogated by methylation-based clocks. Given this role in developmental progression, and prior literature reporting significantly decelerated epigenetic aging in the blood of Kabuki syndrome patients^29,30^, we hypothesized that KMT2D mutant organoids might display epigenetic signatures reflecting a failure to mature normally. Mutant and WT isogenic lines were profiled in biological triplicate at six months of culture. PhenoAge values were consistently more negative in KMT2D mutants than in WT organoids, with minimal overlap between groups. Additionally, profiling with a separate model clock trained on human prenatal brain tissue^47^ showed trends in the same direction, though they did not reach significance due to high variance (**Figure 3A**). In contrast to the accelerated epigenetic aging observed in the SPC2 model, KMT2D-mutant organoids remain epigenetically younger than their wild-type counterparts, consistent with developmental arrest and KMT2D’s established role in stabilizing lineage identity during neurodevelopment. The direction of this effect is concordant with the negative epigenetic age acceleration reported in Kabuki syndrome patient blood^29^, although the magnitudes are not comparable across tissues, clock models and experimental systems.

**Figure 3.**
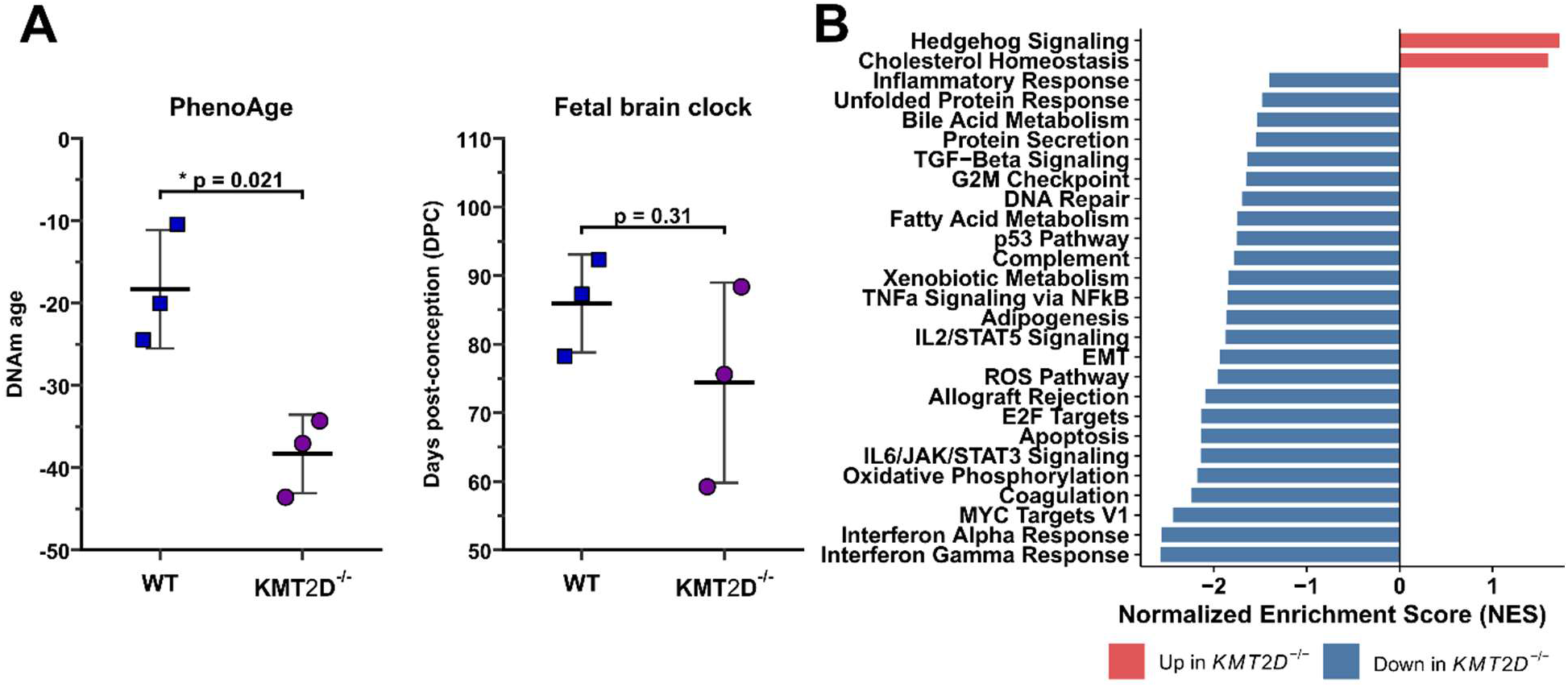
KMT2D mutant brain organoids show epigenetic and transcriptional signatures consistent with developmental arrest. **(A)** Predicted PhenoAge and fetal brain age (measured in days post conception) in WT and KMT2D^-/-^ cortical brain organoids at six months of culture. Individual data points are shown with mean ± SD (n=3 per group). Unpaired t-test with Welch’s correction. (p<0.05.) **(B)** Gene set enrichment analysis (GSEA) of bulk RNA-seq data from KMT2D^-/-^ versus WT brain organoids. Bars represent normalized enrichment score (NES) for each significantly enriched term (FDR < 0.05). Red bars indicate positive enrichment (upregulated in KMT2D^-/-^); blue bars indicate negative enrichment (downregulated in KMT2D^-/-^). Analysis performed against the MSigDB Hallmark gene set database.

### KMT2D loss produces a distinct developmental epigenetic and transcriptional state

KMT2D mutant brain organoids displayed a methylation pattern distinct from both extended culture and the SPC2 disease model. The highly methylated CpG peak was reduced relative to WT organoids, but the corresponding redistribution again occurred across intermediate methylation values (**Supplemental Figure S1C**). The KMT2D comparison yielded a considerably smaller gene list (207 unique DMR-proximal genes), with no terms surviving multiple testing correction (**Supplemental Table 3**). Hedgehog Signaling and Interferon Gamma Response appeared nominally, consistent with the other comparisons, but statistical power was limited.

Transcriptional analysis provided a clearer view of the underlying biological state. GSEA of the KMT2D transcriptional data identified positive enrichment of Hedgehog Signaling and Cholesterol Homeostasis in KMT2D mutants, whereas a broad set of programs spanning immune and inflammatory responses, stress and DNA damage responses, metabolism, and proliferation were negatively enriched (**Figure 3B; Supplemental Table 4**). Many of these suppressed programs overlapped with pathways enriched in the passaging and SPC2 methylation comparisons (**Supplemental Table 7**).

This inverse relationship suggests that biological programs activated during organoid maturation, extended culture, or disease-associated stress remain transcriptionally suppressed following KMT2D loss. Together with the more negative PhenoAge values observed in mutant organoids, these findings support a developmentally stalled epigenetic state rather than accelerated aging, and reflect the known role of KMT2D in activating developmental enhancer programs and stabilizing lineage identity during neurodevelopment^46,48^.

### Sickle cell disease accelerates epigenetic age in iPSC-derived erythroid progenitors

To determine whether disease-associated modulation of epigenetic age extends beyond three-dimensional organoid systems, we examined a two-dimensional iPSC-derived erythroid model of sickle cell disease (SCD). Although SCD arises from a single mutation in the β-globin gene, it is a multisystem disorder characterized by chronic hemolysis, vaso-occlusion, endothelial dysfunction, oxidative stress, and persistent inflammation^49,50^. These processes contribute to progressive injury across the hematopoietic, vascular, pulmonary, renal, hepatic, neurological, and musculoskeletal systems, leading to premature accumulation of age-associated morbidities and accelerated biological aging *in vivo*^51–53^. These observations suggest that chronic disease burden in SCD is reflected in blood-based epigenetic aging measures and provide a clinically relevant context for evaluating whether disease-associated aging signatures can re-emerge in iPSC-derived erythroid cells.

Erythroid progenitors were differentiated from an SCD patient-derived iPSC line and an unaffected control line and profiled at days 20 and 25. At day 25, predicted epigenetic age was significantly elevated in SCD progenitors across the Horvath pan-tissue clock, the skin & blood clock and PhenoAge, as well as the Hannum clock (another blood-based predictor). (**Figure 4A**). Although the absolute differences were less dramatic, their concordant direction across these independently trained clock models supports the presence of a disease-associated epigenetic age shift in a non-organoid iPSC-derived system. Because the SCD and control lines were derived from different donors rather than forming an isogenic pair, however, the magnitude of this effect should be interpreted cautiously.

**Figure 4.**
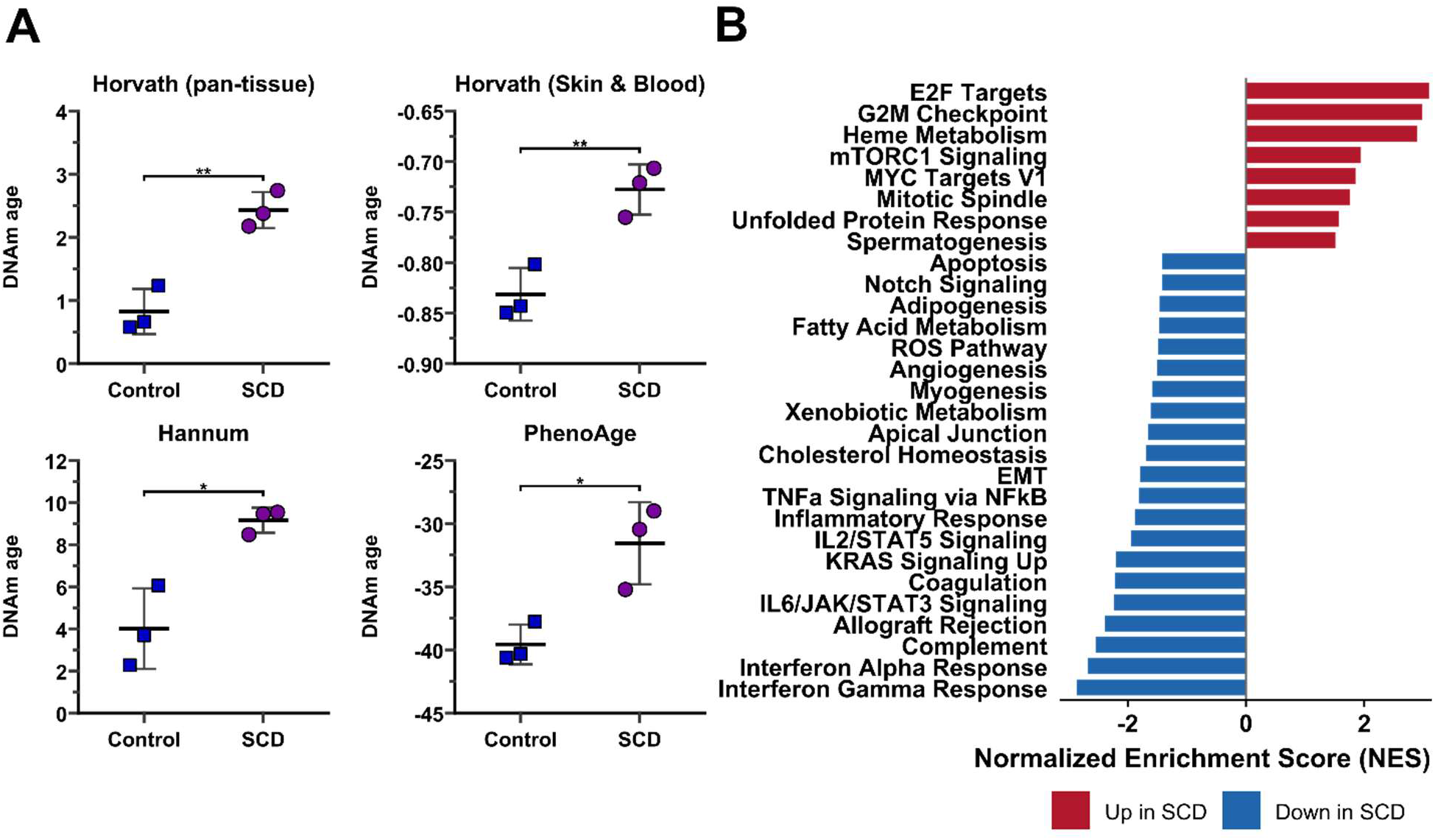
Sickle cell disease iPSC-derived erythroid progenitors show elevated epigenetic age and a distinct transcriptional state. **(A)** Predicted epigenetic age in control and sickle cell disease iPSC-derived erythroid progenitors at day 25 of differentiation, estimated with four blood-based DNA methylation clocks (Horvath pan-tissue, Horvath skin & blood, Hannum, and PhenoAge). Bar graphs show mean ± SD (n=3 per group). Unpaired t-tests with Welch’s correction (p < 0.005). **(B)** Gene set enrichment analysis (GSEA) of bulk RNA-seq data from sickle cell disease versus control erythroid progenitors at day 25. Bars represent normalized enrichment score (NES) for each significantly enriched term (FDR < 0.05; 29 terms total). Red bars indicate positive enrichment (upregulated in SCD); blue bars indicate negative enrichment.

To define the transcriptional state accompanying this epigenetic age increase, we performed preranked gene set enrichment analysis of bulk RNA-sequencing data. At day 25, SCD progenitors showed positive enrichment of proliferative and biosynthetic programs, including E2F Targets, G2M Checkpoint, MYC Targets, mTORC1 Signaling, and heme metabolism (**Figure 4B; Supplemental Table 5**). This profile is consistent with the elevated erythropoietic demand and progenitor proliferation associated with SCD. Because proliferative history and cell division can contribute to methylation changes captured by epigenetic clocks, increased proliferative activity may partly underlie the elevated age estimates observed in these cells.

In contrast, multiple immune and inflammatory pathways were negatively enriched at day 25, despite the well-established systemic inflammatory state of SCD. This pattern was stage-dependent rather than uniform across differentiation: at day 20, TNFα signaling, Inflammatory Response, and IL2-STAT5 Signaling were positively enriched in SCD progenitors before reversing at day 25 (**Supplemental Tables 5 and 6**). These temporal changes suggest that inflammatory pathway activity evolves during erythroid maturation and that a single differentiation stage may not capture the full disease-associated inflammatory trajectory.

Together, these findings extend disease-associated epigenetic age modulation beyond organoid systems to a two-dimensional erythroid model. The persistence of an elevated epigenetic-age signal following somatic-cell reprogramming and redifferentiation is particularly notable because the systemic inflammatory, vascular, and organ-level environment experienced by individuals with SCD is absent from this culture system. Although donor-background effects cannot be excluded, the results raise the possibility that SCD-associated genetic and cell-intrinsic erythroid abnormalities are sufficient to re-establish at least a component of the accelerated epigenetic-aging phenotype observed in patient blood.

## Discussion

In this study, we demonstrate that iPSC-derived organoids accumulate epigenetic age in a time-dependent and reproducible manner, and that disease-associated mutations can modulate this trajectory. Alveolospheres progressively acquired predicted epigenetic ages in some models of nearly 30 years over extended culture, accompanied by a global shift toward hypomethylation that mirrors canonical aging-associated epigenetic drift. Isogenic disease models carrying an SPC2 mutation (alveolospheres) or a KMT2D mutation (cortical brain organoids) displayed distinct epigenetic age profiles relative to their controls, with divergent clock model responses reflecting the underlying biology of each condition, and this modulation extended to non-organoid, two-dimensional erythroid model of sickle cell disease. Together, these findings suggest that iPSC-derived organoids represent a tractable platform for the study of epigenetic aging, and that the combination of extended culture and disease-relevant genetic context can restore substantial epigenetic variation to reprogrammed cell systems.

A central challenge in iPSC-based aging research is that reprogramming resets the epigenetic landscape of these cells, effectively erasing the molecular signatures of age that may motivate the use of specific lines in the first place. In this context, the progressive epigenetic age accumulation observed across multiple organoid systems across a large number of models is particularly notable. The low variance across three independent biological replicates at each alveolosphere timepoint argues strongly that this accumulation is a robust and reproducible feature of the organoid system rather than technical noise. Why might organoids accumulate significant epigenetic age relative to monolayer cultures? While it is possible that the three-dimensional structure and cellular composition of these organoids more faithfully recapitulate the *in-vivo* environment of aging tissues, it is also possible that the deeper level of cellular complexity in these systems recapitulates elements of development which are fundamentally intertwined and linked to the epigenetic changes that occur with aging.

A recurring theme across our dataset is the differing sensitivity of clock models to disease conditions. In the healthy alveolosphere time course, the Horvath pan-tissue clock and PhenoAge behaved concordantly, both tracking the accumulation of epigenetic age with passaging. In the SPC2 and KMT2D disease models, however, disease mutations predominantly manifested in PhenoAge readouts. Rather than treating this as a limitation, we argue that it is itself informative. The Horvath clock was designed to track chronological age across tissue types, while PhenoAge was calibrated to predict biological age in relation to morbidity and mortality risk. These distinct training objectives mean that the two models are sensitive to different features of the methylation landscape, and the selective responsiveness of PhenoAge to disease mutations may reflect the fact that these mutations alter the epigenetic landscape in ways relevant to biological aging rather than chronological aging.

Perhaps the most conceptually interesting finding of this study is the directional contrast between two of our disease models. The SPC2 mutation, which causes proteostatic stress, inflammation, impaired autophagy, and metabolic dysregulation (overlapping substantially with hallmarks of aging), is associated with elevated PhenoAge relative to corrected controls. The KMT2D mutation, which disrupts developmental enhancer activation, and impairs lineage commitment, is associated with more negative PhenoAge relative to wild type controls. In both cases, the direction of the epigenetic age shift is consistent with the known biology of the mutation: one that resembles accelerated aging, and one that resembles developmental arrest.

The global methylation analysis provides orthogonal evidence for the biological significance of the epigenetic age accumulation we observe. The progressive shift toward hypomethylation across alveolosphere timepoints recapitulates an established epigenetic hallmark of *in vivo* aging, suggesting that the underlying methylation changes are not confined to the small number of CpG sites interrogated by clock models but are genome-wide in scope.

At the pathway level, the DMR-based overrepresentation analysis (ORA) and transcriptional GSEA converge on several themes. The passaging comparison is enriched for immune, inflammatory, and EMT-associated programs consistent with inflammaging and age-related fibrotic processes. The SPC2 comparison shares several of these features but additionally shows significant enrichment for Inflammatory Response, consistent with the known disease phenotype. The KMT2D transcriptional data instead reveals broad downregulation of stress response, immune, metabolic, and proliferative programs, consistent with a developmentally arrested rather than aged epigenetic state. That many programs transcriptionally suppressed in KMT2D mutants are epigenetically enriched in the passaging and SPC2 comparisons supports the interpretation that these programs are normally activated as cells mature and age, and that KMT2D loss prevents this activation. Additionally, the consistent enrichment of Hedgehog Signaling across organoid models at varying levels of statistical confidence is an intriguing cross-condition finding. Hedgehog plays fundamental roles in both lung and brain development, and has been implicated in aging-related processes including tissue repair, stem cell maintenance, and fibrosis. Its appearance across tissue types and disease contexts raises the possibility that Hedgehog pathway loci are susceptible to epigenetic remodeling in iPSC-derived organoid systems, though the mechanistic basis for this sensitivity remains to be determined.

Finally, our sickle cell erythroid model indicates that disease-associated modulation of epigenetic age is not confined to organoid systems. SCD progenitors displayed significantly elevated epigenetic age relative to controls across both clock models. This suggests that the capacity of disease state to modulate epigenetic age in iPSC-derived cells is a broader phenomenon than the organoid setting in which we characterize it here.

The findings presented here have several implications for the use of iPSC-derived systems in aging research. If epigenetic age can be reliably induced in organoid systems, it facilitates the study of aging-relevant epigenetic variation in patient-specific genetic backgrounds that would otherwise be inaccessible following reprogramming, including rare populations such as centenarians and individuals with exceptional longevity, where the epigenetic basis of resilience remains to be fully understood. The bidirectional modulation of epigenetic age by disease mutations further suggests that organoid systems may be useful for dissecting how specific genetic variants interact with the aging epigenome, complementing existing approaches based on transcriptional or proteomic profiling. Realizing this potential will require more systematic characterization of the relationship between clock readouts and functional aging phenotypes in organoid systems, and continued development of clock models suited to diverse tissue contexts beyond blood. The present study provides a foundation and motivation for pursuing these goals.

## Limitations of study

Several limitations of this study should be noted. Firstly, the tissue-context limitation of PhenoAge warrants acknowledgment. While this clock was developed using blood-derived methylation data, Levine et al. demonstrated that PhenoAge correlates strongly with chronological age across all tissues and cell types tested^19^, suggesting that its underlying CpG sites capture a broad array of aging-associated methylation changes. That said, absolute predicted ages in non-blood contexts cannot be directly interpreted as biological ages in a physiological sense, and the performance of blood-trained clocks in non-blood tissues is an acknowledged limitation of the field. We argue, however, that in the context of isogenic comparisons, where the only variable is the presence or absence of a disease mutation, against a matched differentiation protocol and genetic background, the relative difference between groups is the meaningful quantity rather than the absolute value. The consistent and large-magnitude differences we observe between mutant and corrected lines, which substantially exceed inter-replicate variance in both groups, support the interpretation that PhenoAge is capturing real and reproducible epigenetic variation rather than tissue-context noise. Secondly, the absence of transcriptional data for the alveolosphere conditions precludes direct comparison of transcriptional and epigenetic changes across tissue types and constrains the interpretation of the ORA findings to the methylation level alone. The KMT2D disease comparison is also limited to a single timepoint, limiting our ability to assess how the KMT2D epigenetic phenotype evolves over time. Thirdly, In contrast to the isogenic SPC2 and KMT2D comparisons, the sickle cell disease model compares a patient-derived line against an unrelated healthy control line rather than an isogenic pair. Consequently, the transcriptional and epigenetic differences observed cannot be fully disentangled from differences in donor genetic background, and the magnitude of the epigenetic age difference in this model should be interpreted with corresponding caution. Lastly, sample sizes are relatively small across specific disease comparisons, with n=2 for SPC2 mutant groups at both timepoints in the methylation data, and n=2 for KMT2D bulk RNAseq samples, limiting statistical power. Conclusions from these groups are supported primarily by the magnitude and consistency of the observed effects rather than formal statistical inference.

## Methods

### iPSC-directed differentiation into alveolar epithelial type 2 cells (iAT2s) and iAT2 maintenance

In this study, we used our previously published SPC2 iPSC line clones SPC2-ST-B2 (SFTPC^tdT/WT^; “SFTPC corrected”) and SPC2-ST-C11 (SFTPC^I73T/tdT^; “SFTPC^I73T^ mutant”)^42^. As previously described^42^, these clones carry an SFTPC^tdTomato^ reporter. iPSCs used in this study demonstrated a normal karyotype when analyzed by G-banding and/or array Comparative Genomic Hybridization (Cell Line Genetics). We used our previously published directed differentiation protocol^34,42^ to generate SFTPC^I73T^ mutant and corrected iAT2s. In brief, the STEMdiff Definitive Endoderm Kit (Stemcell Technologies, #05110) was used for differentiation into endoderm. On day 3 cells were dissociated using Gentle Cell Dissociation Reagent (StemCell Technologies, #07174), passaged onto 6-well tissue culture plates coated with growth factor reduced Matrigel (Corning 354277) and cultured in ‘‘DS/SB’’ anteriorization medium, consisting of complete serum-free differentiation medium (cSFDM) base as previously described^34,35^, supplemented with 2 μM Dorsomorphin (“DS”, Fisher Scientific, #NC0275327) and 10 μM SB431542 (“SB”, Sigma, #S4317). After 3 days of culture in “DS/SB” medium, cells were cultured in cSFDM supplemented with 3 μM CHIR99021, 10 ng/ml rhBMP4 (R&D Systems, #314-BP), and 100 nM Retinoic Acid (Sigma, #R2625). On day 15-16 of differentiation, cells were sorted for lung progenitors based on CD47^hi^/CD26^neg^ gating^54^. Sorted cells were resuspended in 50 µl droplets of 3D Matrigel (Corning, #354230) at 400 cells/μl and cultured in alveolar differentiation medium “CK+DCI”, consisting of cSFDM base supplemented with 3 μM CHIR99021 (Fisher Scientific, 44-235-0), 10 ng/ml rhKGF (Fisher Scientific, #251KG050), 50 nM Dexamethasone (Sigma, #D4902), 0.1 mM 8-bromoadenosine 3’,5’ cyclic monophosphate sodium salt (Sigma, # B7880), and 0.1 mM 3-isobutyl-1-methylxanthine (IBMX, Sigma, # I5879). CK+DCI medium was supplemented with 10 mM Y-27632 (Tocris, #1254) for the first 48 h after sorting, and cells were fed with fresh CK+DCI medium every 48-72 h. Epithelial spheres were passaged between day 28 and 37 of differentiation, and a brief period of CHIR withdrawal was subsequently performed to achieve iAT2 maturation as previously described^35^. SFTPC^tdTomato^+ cells were then purified by FACS sorting and plated at 400 cells/μl in 3D Matrigel (Corning, #354230). iAT2s were subsequently maintained through serial passaging and re-plating into 3D Matrigel droplets at a density of 400 cells/μl every 10 days. For passaging, cells were dissociated into single cells with 2 mg/ml Dispase (Gibco, #17105-041) for 1 h at 37 °C, followed by incubation in 0.05 % Trypsin (Gibco, #25300062) at 37 °C for 12 mins. iAT2 purity was assessed by flow cytometry for SFTPC^tdTomato^ expression. Paired SFTPC^I73T^ mutant and corrected iAT2s at the indicated days of differentiation were used in this study.

### Brain organoid generation (KMT2D model)

WTC11 (WT) and KMT2D^-/-^ hiPSC lines were provided by the Serrano laboratory (Boston University, Center for Regenerative Medicine). The KMT2D^-/-^ line (clone KMT2D.10) was generated in the WTC11 background by CRISPR/Cas9-mediated introduction of a premature stop codon in exon 2 of KMT2D, as previously described^46^. Cerebral organoids were generated and maintained according to the telencephalic protocol described in Ullah et al. 2025^55^, as implemented by Golden et al. 2025^46^. Briefly, hiPSCs were subjected to dual SMAD inhibition in N2/B27 medium, followed by neural rosette expansion in EGF/FGF-containing medium, and subsequent maturation in suspension culture. Organoids were harvested for DNA extraction at approximately six months of culture. Individual organoids were dissociated in papain solution and pelleted; DNA was extracted from the resulting cell pellet for methylation array profiling. No live cell selection was performed prior to extraction.

### Generation of erythroid progenitors (SCD model)

Patient-specific SCD iPSC line SA170, established as part of a comprehensive SCD iPSC library^56^, and healthy control iPSC line BU6 were differentiated into hematopoietic stem and progenitor cells (HSPCs) and subsequently specified towards the erythroid lineage as previously described^57–60^. In brief, iPSCs were plated on Matrigel (Corning Matrigel hESC-qualified Matrix; #356234) at ∼10% confluency and in mTeSR 48 hours prior to differentiation. iPSCs were then specified towards hemogenic endothelium and HSPCs budding off into the medium were collected on day 13, spun down and resuspended in erythroid specification media (SS1). On day 20, cultures consisting of early-intermediate erythroid progenitors were spun down and replated in erythroid maturation medium (SS2) for further maturation. iPSC-derived erythroid progenitors were collected on day 20 and day 25 for profiling in this work. SS1 media consists of: StemSpan SFEM II (StemCell Technologies; #09655), 2mM l-glutamine (Gibco; #25030164), 100ug/mL Primocin (InvivoGen; #ant-pm-2), 100ng/ml hSCF (R&D Systems; #255-SC), 40ng/mL recombinant human IGF-1 protein (IGF-1) (R&D Systems; 291-G1), 0.5μM Dexamethasone (Sigma-Aldrich; #D4902), 0.5 U/ml human erythropoietin (EPO) (Henry Schein; #1265340). SS2 media corresponds to SS1 media without hSCF, IGF-1 and Dexamethasone. All cells were cultured at 37°C in normoxic, 5% CO2 conditions.

### iPSC reprogramming time course

Reprogramming of PBMCs from a 54-year-old donor to induced pluripotency was performed using Sendai virus-mediated delivery of reprogramming factors, as implemented in Dowrey et al. (2024)^61^. Briefly, thawed PBMCs were expanded in erythroblast expansion medium for 9 days, after which cells were transduced using the Invitrogen CytoTune-iPS 2.0 Sendai Reprogramming Kit (Thermo Fisher A16517). The following day, cells were transferred onto Matrigel-coated wells and gradually transitioned into ReproTeSR medium (STEMCELL Technologies 05926) over the first 5 days post-transduction (ReproTeSR added on Days 3 and 5 without aspirating the existing expansion medium; from Day 5 onward cells were maintained fully in ReproTeSR). Once colonies were sufficiently large, individual colonies were divided by a cross-hatching method and passaged onto Matrigel-coated wells to establish clonal cultures. To capture epigenetic age across the reprogramming process, cells were collected at Day 0 (pre-transduction), Day 1 (post-transduction), and Days 3, 9, and 13, and clonal cultures were subsequently collected at passage 1 and passage 3.

### DNA methylation profiling

Genomic DNA from our samples was extracted using the Qiagen AllPrep DNA/RNA Kit (80204). Extraction was performed according to the manufacturer’s protocol in each case. Purified DNA was submitted to TruDiagnostic for DNA methylation profiling on the Illumina Infinium MethylationEPIC v2.0 BeadChip array. Resulting IDAT files were processed as described under “Data processing and normalization.”

### RNAseq profiling

Total RNA was extracted using the Qiagen AllPrep DNA/RNA Kit (80204) according to the manufacturer’s protocol. Read alignment and quantification are described under “RNA-seq processing and differential expression analysis.”

### Data processing and normalization

DNA methylation profiling was performed using the Illumina HumanMethylationEPIC v2 BeadChip array. Raw methylation data were imported from EPIC v2 array IDAT files using the minfi R package (v1.54.1) and initial quality control was performed to compute mean detection p-values for each CpG site and sample, excluding those with mean detection p-values greater than 0.05 from further analysis. Signal intensities were normalized using single-sample Noob (ssNoob) normalization via the preprocessNoob function in minfi, using the single dye bias correction method. Beta values were collapsed to probe family identifiers using the betasCollapseToPfx function from the SeSAMe R package (v1.26.0), retaining the mean value across replicate probes. Missing CpG values were imputed using reference data from the sesameData package. Following quality control and normalization, the resulting beta values were used as input for epigenetic age estimation across a panel of established methylation clock models. Final predictions are reported as estimated ages in years or unitless age-associated scores depending on the design of each clock model.

### RNA-seq processing and differential expression analysis

Raw reads were aligned to the hg38 reference genome using STAR(v2.7.10b), with gene-level read counts generated using featureCounts (v2.0.3) against the GENCODE v39 annotation. Multimapping and multi-overlapping reads were excluded. Differential expression analysis was performed using DESeq2, comparing KMT2D^-/-^ samples (n=2) versus WT samples (n=2). Low-count genes were filtered prior to analysis, retaining only genes with a minimum of 10 counts in at least 2 samples. For gene set enrichment analysis, genes were ranked by the DESeq2 Wald statistic, which incorporates both the magnitude and statistical significance of differential expression. Preranked GSEA was performed using the fgsea R package against the MSigDB Hallmark gene set database. Gene sets with FDR < 0.05 were considered statistically significant.

### Differentially methylated region analysis

DMR analysis was performed using the DMRcate R package. Prior to DMR calling, probes near single nucleotide polymorphisms were removed using the rmSNPandCH function. Beta values were annotated using the IlluminaHumanMethylationEPICv2anno.20a1.hg38 annotation package. CpG-level statistics were computed using cpg.annotate with the following parameters: arraytype = “EPICv2”, epicv2Filter = “mean”, epicv2Remap = TRUE, analysis.type = “differential”, fdr = 0.001. DMRs were called using dmrcate with lambda = 1000 and C = 2. DMR coordinates were extracted and annotated to the hg38 genome using extractRanges.

### DMR-proximal gene list derivation and filtering

Genomic regions identified as significant DMRs by DMRcate were annotated to overlapping genes using the extractRanges function against the hg38 genome annotation. Genes whose gene body or promoter region overlapped a DMR were retained; non-coding and antisense genes were then removed prior to pathway analysis.

### Global methylation distribution analysis

Genome-wide beta value distributions were visualized as smoothed density curves for each sample using the density function in R. Distributions were plotted across all autosomal CpG probes passing quality control thresholds to assess global methylation patterns across conditions and timepoints.

### Over-representation analysis and GSEA

Over-representation analysis (ORA) of DMR-proximal gene lists was performed using the enrichR R package against the MSigDB Hallmark gene set database. ORA was used in preference to GSEA for our DMR-derived gene lists, which lack the inherent ranking or directionality required for GSEA. The default EnrichR background gene set was used. Enriched terms with FDR < 0.05 were considered statistically significant; terms with nominal p < 0.05 but FDR ≥ 0.05 are reported as nominally enriched. Preranked gene set enrichment analysis was performed using the GSEA desktop application against the MSigDB Hallmark gene set collection (v2024.1.Hs), with genes ranked by the DESeq2 Wald statistic. Ensembl gene identifiers were collapsed to gene symbols using the MSigDB Human Ensembl Gene ID chip (v2024.1.Hs) with the Abs_max_of_probes collapse mode.

### Statistical analysis

All graphical statistical analyses were performed using GraphPad Prism (version 8.0.2). Epigenetic age across three timepoints in the alveolosphere time course was assessed by two-way ANOVA with Tukey’s multiple comparisons correction, with clock model and timepoint as factors. Comparisons of PhenoAge between SPC2 mutant and corrected alveolospheres at D97 and D209 were performed using unpaired t-tests with Welch’s correction, with Holm-Šídák adjustment applied across the two timepoint comparisons (corrected: n=3; SPC2 mutant: n=2). Comparison of PhenoAge between KMT2D mutant and WT brain organoids was performed using an unpaired t-test with Welch’s correction (n=3 per group). Epigenetic age comparisons between D25 SCD and control erythroid progenitors were performed using unpaired t-tests with Welch’s correction (n = 3 per group). Alpha was set at 0.05 for all comparisons. Data are presented as mean ± standard deviation. Normality was assumed given the continuous nature of the data and limited sample sizes precluding formal normality testing.

## Acknowledgments

We thank and acknowledge the Kotton Laboratory in the Center for Regenerative Medicine for generation of our iAT2 cells (clones SPC2-ST-B2 and SPC2-ST-C11) used in this study.

## Author contributions

Conceptualization: NJS, TWD, GJM

Methodology: NJS, TWD, SC, EM, RG, DJ, CSG, BP, ML, MAS, KA, KV, GJM

Investigation: NJS, GJM Writing: NJS

Review and editing: NJS, TWD, MAS, KA, KV, GJM

## Data and materials availability

Data and materials are available on GEO under repository number GSE346747.

