## Supplemental Figures S1 for "Disease-associated mutations directionally modulate epigenetic age in iPSC-derived tissues"

**Supplementary Figure 1. Global methylation distributions across organoid conditions.**

Genome-wide beta value density distributions for **(A)** alveolosphere organoids at D97, D209, and D360+ culture timepoints, **(B)** SPC2 mutant versus corrected isogenic alveolosphere organoids at D97, and **(C)** KMT2D^-/-^ mutant versus WT brain organoids at six months. Each line represents an individual sample. Beta values approaching 0 indicate unmethylated CpGs; beta values approaching 1 indicate methylated CpGs.

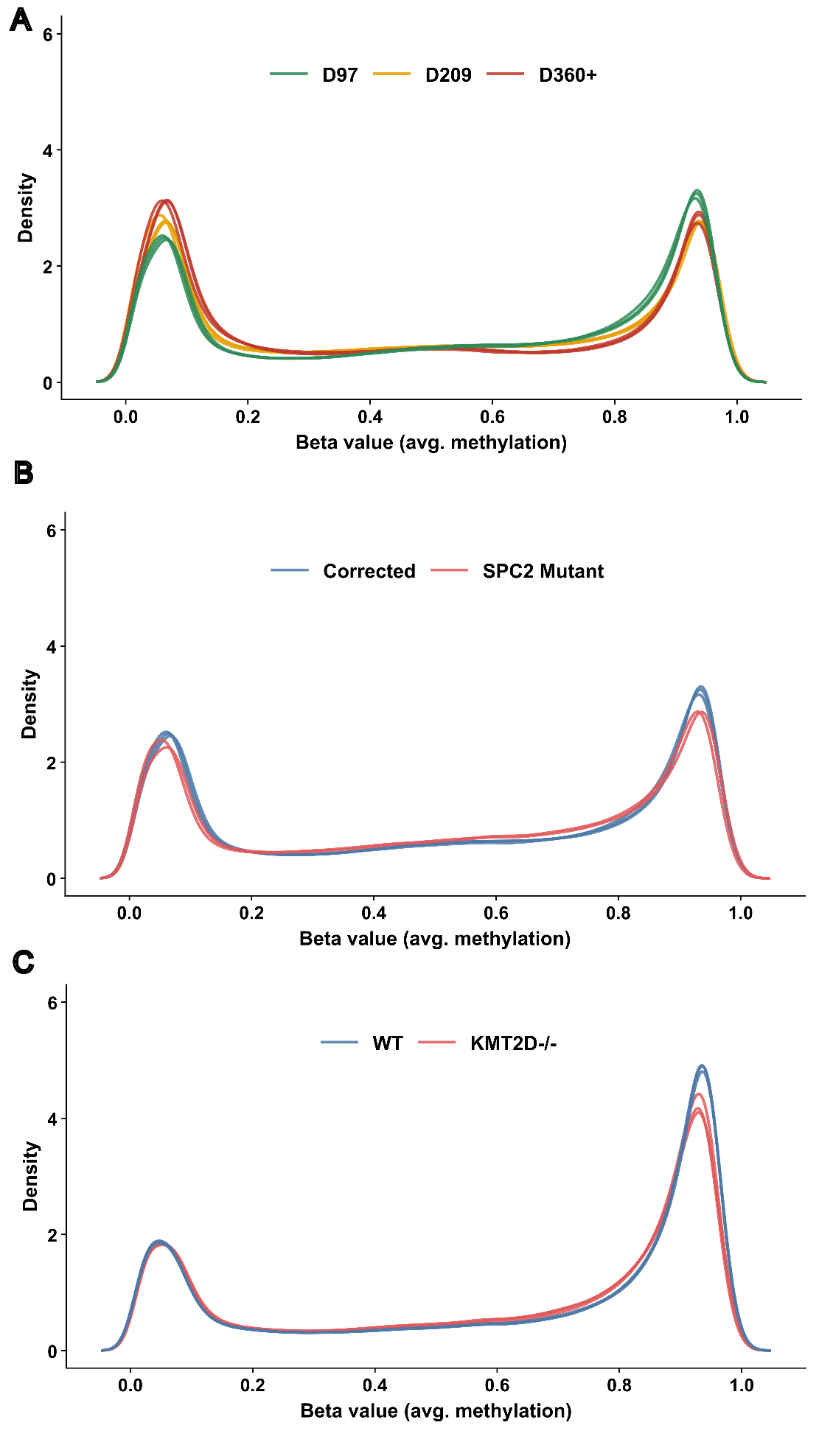
